# A Cell Proliferation and Inflammatory Signature is Induced by *Lawsonia intracellularis* Infection in Swine

**DOI:** 10.1101/384230

**Authors:** Fernando L. Leite, Juan E. Abrahante, Erika Vasquez, Fabio Vannucci, Connie J. Gebhart, Nathan Winkelman, Adam Mueller, Jerry Torrison, Zachary Rambo, Richard E. Isaacson

## Abstract

*Lawsonia intracellularis* causes porcine proliferative enteropathy. This is an enteric disease characterized by thickening of the wall of the ileum that leads to decreased growth and diarrhea of animals. In this study, we investigated the host response to *L. intracellularis* infection by performing transcriptomic and pathway analysis of intestinal tissue in groups of infected and non-infected animals at 14, 21 and 28 days post challenge. At the peak of infection, when animals developed most severe lesions, infected animals expressed higher levels of several genes involved in cellular proliferation and inflammation including matrix metalloproteinase-7 (*MMP7*), transglutaminase-2 (*TGM2*) and Oncostatin M (*OSM*). Histomorphology also revealed general features of intestinal inflammation. This study identified important pathways associated with the host response in developing and resolving lesions due to *L. intracellularis* infection.

**Importance:** *Lawsonia intracellularis* is among the most important enteric pathogens of swine and it can also infect other mammalian species. Much is still unknown regarding its pathogenesis and the host response, especially at the site of infection. In this study we uncovered several novel genes and pathways associated with infection. These differentially expressed genes, in addition to histological changes in infected tissue, revealed striking similarities between *L. intracellularis* infection and cellular proliferation mechanisms described in some cancers and inflammatory diseases of the gastrointestinal tract. This research sheds important light into the pathogenesis of *L. intracellularis* and the host response associated with the lesions caused by infection.

## Introduction

*Lawsonia intracellularis* is a Gram-negative, obligate intracellular bacterium that infects enterocytes of swine, primarily in the ileum, and causes a disease called proliferative enteropathy (PE) (1). Infected enterocytes undergo hyperplasia and macroscopic lesions are marked by thickening of the intestinal mucosa (1). PE is endemic in swine herds worldwide (1, 2) and *L. intracellularis* has also been shown to infect horses, hamsters, dogs, and non-human primates among other species (3).

In swine, there are two major clinical forms of disease: proliferative hemorrhagic enteropathy (PHE) and porcine intestinal adenomatosis (PIA). PIA is a persistent but self-limiting disease that occurs in young pigs and can lead to diarrhea, reduced growth, and is commonly a subclinical disease (1). PHE occurs in older finisher pigs, gilts, and sows and is characterized by hemorrhagic diarrhea and often leads to death (1). PIA is the most common form of the disease (3) and was the focus of this study.

There is limited knowledge on the pathogenesis of *L. intracellularis*. This gap in knowledge is partly due to the fastidious nature of the organism, the difficulty growing pure cultures of the organism, and the lack of *in vitro* models that replicate proliferative lesions (3). Similarly, much is still unknown about the mucosal immune response to *L. intracellularis*. Elevated levels of TNF-α and TGF-β have been found in the infected intestinal mucosa (4, 5) along with an antigen-specific IgA response (6). Infiltration of inflammatory cells, particularly neutrophils, is not a primary feature of infection (1), although accumulation of macrophages has been found to coincide with increased lesion severity and antigen load in this disease (7, 8).

The objective of this study was to investigate the porcine host response to *L. intracellularis* at the site of infection to gain a better understanding of the pathogenesis and immune response by correlating the presence and severity of lesions with the differential expression of host genes at several time points using RNAseq and Pathway Analysis. Our results demonstrated that several genes associated with cell proliferation and inflammation are differentially expressed in infected animals, a pattern which is exacerbated with increase in lesion severity.

## Materials and Methods

### Animals and treatments

The animal protocol used was approved by the Swine Services Unlimited, Inc. Institutional Animal Care and Use Committee and all experiments were performed in accordance with relevant guidelines and regulations. Genetically, pigs were from a Landrace cross to a Yorkshire female with a large white sire (Topigs-Norsvin). Two groups of 18 animals were used in this study. One group was infected with *Lawsonia intracellularis* and the other group was not infected. The inoculum used was prepared following a previously described protocol and consisted of a mucosal homogenate which was negative for other bacterial and viral pathogens including porcine reproductive and respiratory syndrome virus (9). Animals were challenged with a dose calculated by qPCR to be of 8.0 x 10^7^ *L. intracellularis* organisms by oral gavage at 7 weeks of age. Groups of 6 pigs per treatment were randomly selected and euthanized at 14, 21 and 28 days post challenge.

### Gross and microscopic pathology evaluation

The entire intestinal tract was examined for lesions characteristic of PE. Each macroscopic lesion was scored as either mild (score of 1), moderate (score of 2) or severe (score of 3). For histopathology and evaluation of microscopic lesions characteristic of PE, a section of ileum proximal to the ileocecal valve was collected from each animal. This section of the intestine was chosen as it has been described as the most consistent site of *L. intracellularis* infection (1, 7). The sections were then stained using hematoxylin and eosin (HE) and used for immunohistochemistry (IHC) by staining using murine anti-L. *intracellularis* specific polyclonal antibody following previously described protocols (10). The presence of *L. intracellularis* specific antigen was measured with a five-grade IHC scoring scale (grade 0 equal to no *L. intracellularis* antigen found in tissue; grade 1 equal to 0-25%, grade 2 equal to 25-50%, grade 3 equal to 50-75% and grade 4, equal to more than 75% enteric crypts containing antigen) (11). The pathologist reading the slides was blinded to which experimental group was being scored. Microscopic lesions in HE sections were measured using a four grade scale representing the distribution of crypt dysplasia: 0 no lesion, 1 for focal lesions, 2 for multifocal lesions, and 3 for diffuse lesion distribution. An animal was considered to have a high level of infection and lesions if the IHC score was 2.5 and above, HE score was 2 and above and there was the presence of gross lesions. Animals with a low level of lesions were defined as those that had IHC and HE scores of one or zero with or without the presence of gross lesions in the group that received the challenge.

### Histomorphometry

To assess differences in crypt and villous morphology due to infection, intestinal sections from animals were evaluated by measuring crypt depth and villous height. For this measurement, images were captured from slides using light microscopy with a 10 x objective and analyzed with ImageJ (12). Twenty intact and well oriented villi were randomly selected along with thirty randomly selected crypts for the measurement of their height and depth, respectively. In instances where twenty villi could not be found, all villi in the image were counted. The average crypt depth and villus height among high lesion, low lesion and non-infected animals was compared using Tukey’s comparison in an analysis of variance (anova) in R (version 3.3.3 (2017-03-06)) with a level of significance set at 0.05.

### Shedding and serologic responses

Antibodies against *L. intracellularis* in serum samples were measured using the immunoperoxidase monolayer assay (IPMA) (13). Fecal shedding of *L. intracellularis* was measured using qPCR (14).

### Differential gene expression and pathway analysis

Immediately following necropsy, a section of ileum 2-3 cm immediately proximal to the ileocecal junction was opened longitudinally and scraped using the edge of a microscope slide. The sample was immediately placed in RNAlater (ThermoFisher) and total RNA was extracted using the RNeasy Plus Universal Mini Kit (Qiagen). RNA quantity and quality were assessed using RiboGreen RNA and Agilent Analysis system, respectively. Samples that passed quality metrics were used to create a cDNA library for sequencing using Illumina Library Creation. Samples from each pig were individually sequenced using the Illumina HiSeq 2500 sequencer using 50 base paired-end reads,20 million reads per sample were obtained. The Illumina sequence files were processed using a pipeline developed by the University of Minnesota Informatics Institute. Briefly, FastQ files were trimmed via trimmomatic and mapping was performed via TopHAT (v2.0.13) using bowtie (v2.2.4.0). The Sus Scrofa 3.0 (susScr3) genome was used and gene annotation was performed using Ensembl from the same genome build. Differentially expressed gene (DEG) analysis was performed using CLC genomics workbench (CLCGWB v 9.0.1, Qiagen Bioinformatics), and EdgeR. EdgeR has been described to have superior specificity and sensitivity as well as good control of false positive errors when compared to other methods to detect DEG (15). Ingenuity^®^ Pathway Analysis (IP A®, QIAGEN) was used for pathway analysis.

Comparisons were made between infected and non-infected animals at each timepoint post infection as well as between animals with high (severe and diffuse) and low-level lesions. At 21 dpi, animals with high lesions (1381 and 97) were compared to the animals with low lesions (144, 173 and192) (table 1). RNA could not be extracted from the intestinal sample of animal 297, which had high lesions. At 28 dpi, high lesion animals were 1385, 189 and 1386 compared to animals 94, 197 and 194 which had low lesions.

**Table 1.** Measures of infection by *L. intracellularis* at different times post infection. Immunohistochemistry, microscopic and gross lesion scores, PCR Ct value and serum antibody titer.

| 14 Dpi |  |  |  |  |  |
| --- | --- | --- | --- | --- | --- |
| Animal ID | IHC | HE | GL | PCR | IPMA |
| 139 | 1 | 1 | 0 | 20.02 | - |
| 99 | 1 | 1 | 0 | 26.33 | 1:30 |
| 148 | 1 | 1 | 0 | 29.44 | - |
| 77 | 1 | 1 | 0 | 34.97 | 1:30 |
| 188 | 1 | 1 | 0 | 26.89 | 1:30 |
| 87 | 1 | 1 | 0 | 27.54 | 1:30 |
| Average | 1.00 | 1.00 | 0.00 | 27.53 | 1:20 |
| 21 Dpi |  |  |  |  |  |
| Animal ID | IHC | HE | GL | PCR | IPMA |
| 144 | 1 | 1 | 0 | 27.11 | 1:480 |
| 173 | 1 | 1 | 0 | 25.51 | 1:240 |
| 192 | 1 | 1 | 1 | 26.3 | - |
| 297 | 3* | 3* | 1 | 27.08 | 1:240 |
| 1381 | 4* | 3* | 3 | 17.33 | 1:240 |
| 97 | 3* | 3* | 1 | 25.04 | 1:240 |
| Average | 2.17 | 2.00 | 1.00 | 24.73 | 1:240 |
| 28 Dpi |  |  |  |  |  |
| Animal ID | IHC | HE | GL | PCR | IPMA |
| 94 | 0 | 1 | 1 | - | 1:240 |
| 197 | 0 | 0 | 1 | - | - |
| 194 | 1 | 1 | 1 | 32.52 | 1:1920 |
| 1386 | 2.5* | 3* | 2 | 26.61 | 1:1920 |
| 1385 | 2.5* | 2* | 1 | 28.62 | 1:480 |
| 189 | 4* | 3* | 2 | 23.61 | 1:960 |
| Average | 1.67 | 1.67 | 1.33 | 27.84 | 1:920 |

dpi= days post infection; IHC= immunohistochemistry score of *L. intracellularis* antigen in tissue; HE= Haemotoxylin and Eosin stain score of microscopic lesions; GL= Gross lesion score; PCR= Ct value; IPMA= serum antibody titer; Neg= result negative; \*= high lesion.

### Immunohistochemistry of matrix metalloproteinase-7

To measure the presence and distribution of metalloproteinase-7 (MMP-7), which is encoded by the gene *MMP7*, IHC was performed on paraffin embedded tissue collected at necropsy. IHC of metalloproteinase-7 was performed using a monoclonal antibody against MMP-7 following an adaptation of a previously described protocol (16), using the primary monoclonal antibody against MMP-7 at a dilution of 1:600. The antibody was obtained from Vanderbilt Antibody and Protein Resource.

## Results

### Gross and microscopic pathology

At 14 days post infection (dpi), all infected animals had IHC and HE scores of one indicating low level infection with minor lesions and no animals had gross lesions (table 1). At 21 dpi, three of 6 animals (297, 1381, 97) had HE lesion scores of three indicating diffuse microscopic lesions and the same three animals also had IHC scores of three and above indicating a high level of *L. intracellularis* present in the tissue. The other infected animals necropsied at this time point (144,173,192) had IHC and HE scores of one indicating minor (low) lesions and one of these animals had mild gross lesions. All three animals at 21dpi with HE and IHC scores above two had gross lesions and one of these animals had severe gross lesions (table 1). At 28 dpi three animals (94, 197, 194) had low lesions with either a negative score or a score of one for IHC and HE. The other three animals at this time point (1386, 1385, 189) had high lesions as measured by IHC and HE scores above 2 and moderate or mild gross lesions (table 1). All non-infected animals did not have microscopic lesions observed by HE or IHC. One animal in the non-infected group at 28 dpi had a gross lesion score of 1 with mild thickening in Peyer’s patches and hyperemic folds (data not shown).

### Shedding and serologic responses

The results of fecal PCR and IMPA serologic assay are shown in table 1. Animals shed more *L. intracellularis* at 21 dpi, when Ct values were the lowest among all three time points (Ct of 24.73 at 21 dpi versus 27.53 at 14dpi and 27.84 at 28dpi). At 28 dpi two animals in the infected group did not shed detectable levels of *L. intracellularis* in feces. In the non-infected group, two animals shed *L. intracellularis* (both with a Ct of 33). Since these animals did not have microscopic lesions measured using HE or IHC and did not have positive antibody responses, they were still included in the study. Antibodies against *L. intracellularis* were not detected in serum samples of any of the animals in the non-infected group. Five of the six animals in the infected group had detectable antibody titers at 21 and 28 dpi. Antibody titers ranged from an average of 1:20 at 14 dpi to 1:920 at 28dpi (table 1).

### Histomorphology

Analysis of crypt and villus morphology revealed that infection affected the intestinal structure, and animals with high lesions responded differently than those with low lesions. Although not statistically significant, at 14 dpi infected animals on average had larger crypts, which peaked at 21dpi in high lesion animals compared to low lesion and non-infected animals. At 28 dpi high lesion and non-infected animals had similar crypt depths (Figure 1a). Villous height was also affected by infection. At 14 dpi, villi height was lower in infected animals (*p*<0.05, Figure 1b). At 21dpi animals in the high lesion group had the shortest villi height, which was significantly shorter than the non-infected group (*p*<0.05, Figure 1b). While crypt depth was similar at 28dpi, villi height was not, and high lesion animals had significantly lower villi height compared to other infected and non-infected animals (*p*<0.05, Figure 1b). Differentially expressed genes between infected and non-infected animals

**Figure 1.**
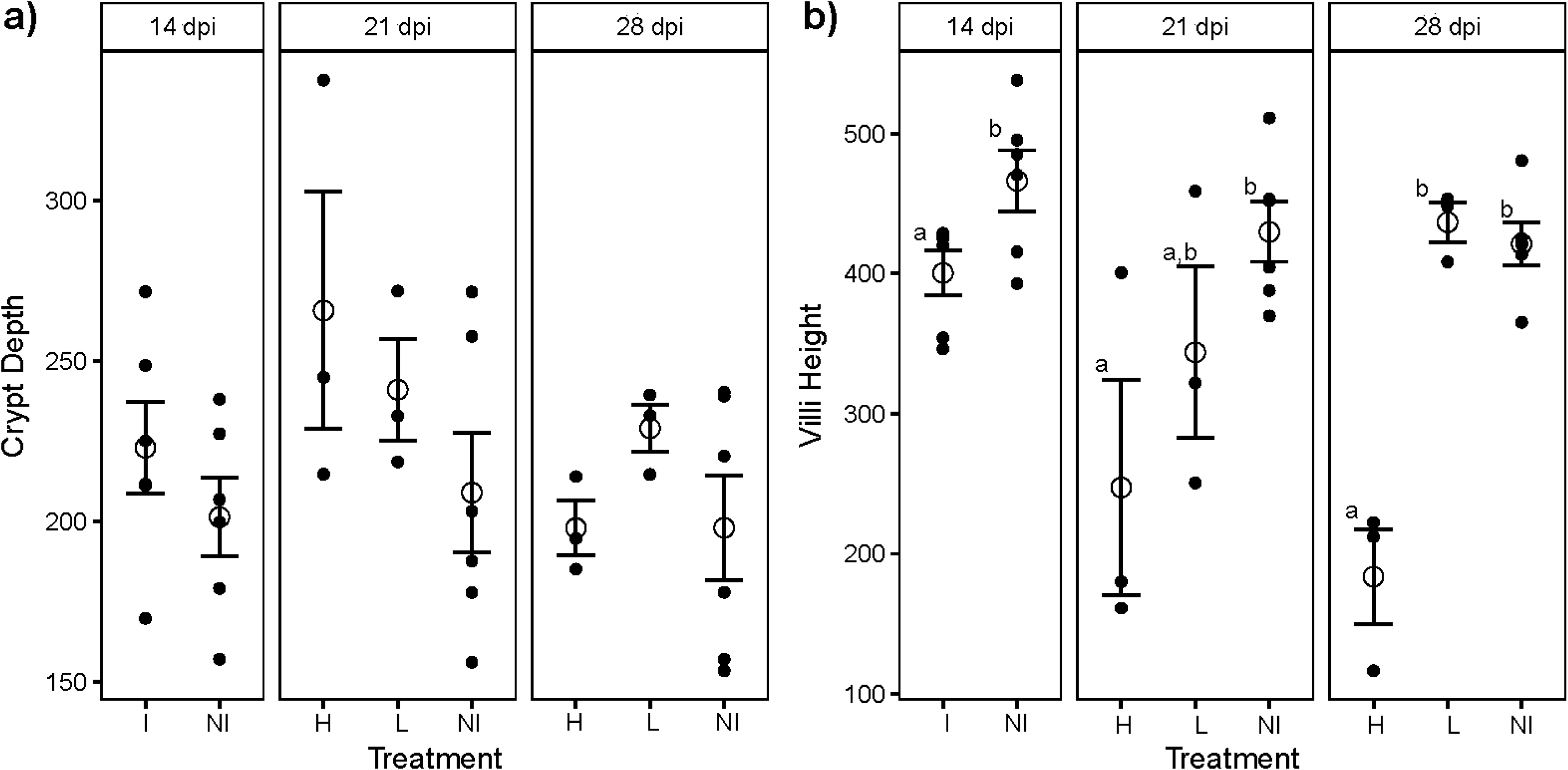
Histomorphology of ileum tissue. a) Crypt depth; b) Villus height. Reported are the means for treatment by day with the standard error. Different letters indicate statistical significance (*p* < 0.05). Error bars represent standard error of the mean, open circles indicate the mean. dpi= days post infection; I= Infected; NI= Non infected; H= High Lesion; L= Low Lesion.

To determine if tissue lesions corresponded to differences in the profile of genes expressed in the ileal samples, RNAseq was employed to compare mRNA profiles between challenged and non-challenged pigs and between pigs with high and low lesions due to *L. intracellularis* infection. When we compared samples from challenged and non-challenged pigs, no altered gene expression profiles were observed at 14 days post challenge. However, at 21 dpi, there were 22 DEGs while at 28 dpi only a single DEG was identified.

Among the 22 DEGs at 21 dpi, were two genes involved in nucleotide metabolism: xanthine dehydrogenase (*XDH*) and cytidine deaminase (*CDA*) (table 2). The gene encoding gamma-aminobutyric acid receptor subunit pi (*GABRP*) was also among the most highly expressed genes. Two matrix metalloproteinase (MMPs) genes were up regulated and these were *MMP7* and *MMP13*.

**Table 2.** Differentially expressed genes identified at 21 days post infection comparing infected to non-infected pigs.

| Ensembl id | Gene Symbol | Fold Change | Corrected *P*-value |
| --- | --- | --- | --- |
| ENSSSCT00000009326.2 | *XDH* | 816.98 | 0.009 |
| ENSSSCT00000016344.2 | *MMP7* | 409.6 | 0.014 |
| ENSSSCT00000026800.1 | *GABRP* | 277.33 | 0.024 |
| ENSSSCT00000014604.2 | - | 198.35 | 3.66E-04 |
| ENSSSCT00000005396.2 | *SERPINB2* | 128.98 | 0.009 |
| ENSSSCT00000014601.3 | *SAA1* | 92.3 | 0.024 |
| ENSSSCT00000004241.1 | *TACSTD2* | 58.54 | 0.016 |
| ENSSSCT00000010382.2 | - | 28.02 | 0.009 |
| ENSSSCT00000029653.1 | - | 25.13 | 0.034 |
| ENSSSCT00000009811.2 | *AMCF-II* | 23.57 | 0.009 |
| ENSSSCT00000000312.2 | - | 23.06 | 0.009 |
| ENSSSCT00000010954.2 | *OSM* | 19.87 | 0.015 |
| ENSSSCT00000011519.2 | *SFRP5* | 18.66 | 0.026 |
| ENSSSCT00000007675.2 | *IDO1* | 17.62 | 0.016 |
| ENSSSCT00000000001.2 | - | 15.66 | 0.024 |
| ENSSSCT00000016351.2 | *MMP13* | 15.04 | 0.021 |
| ENSSSCT00000005162.1 | *DUOXA2* | 13.56 | 0.009 |
| ENSSSCT00000031001.2 | *TGM2* | 11.72 | 0.035 |
| ENSSSCT00000005163.2 | *DUOXA1* | 10.24 | 0.014 |
| ENSSSCT00000026673.2 | *CDA* | 7.3 | 0.013 |
| ENSSSCT00000015915.1 | - | 4.21 | 0.009 |
| ENSSSCT00000010573.2 | *ADAMDEC1* | -4.61 | 0.014 |

Several of the other DEG are known to have roles in inflammatory responses: serum amyloid 1 (SAA1) whose inducible expression is a hallmark of the acute-phase response (17); Oncostatin M (*OSM*) is a cytokine involved in diseases with chronic inflammation (18); secreted frizzled-related protein 5 (*SFRP5*) is an anti-inflammatory adipokine (19); and transglulatminase-2 (*TGM2*) which is a functionally complex protein that is induced by inflammatory mediators (20).

Genes described as being expressed by antigen presenting cells, including macrophages and dendritic cells, were also differentially expressed. These included *SERPINB2, IDO1,* and *ADAMDEC1,* and *AMCF-II* (encoding alveolar macrophage-derived chemotactic factor-II). Other genes up-regulated at 21 days post infection included the tumor-associated calcium signal transducer gene (*TACSTD2*) as well as the genes that code for dual oxidase maturation factor 1 and 2 (*DUOXA2, DUOXA1)*.

The single gene differentially expressed at 28 dpi was unc-5 netrin receptor B (*UNC5B)*. It was expressed 3.77 fold higher in infected animals (p < 0.05).

### Differentially expressed genes between animals with high and low lesions

Because pigs within the challenged group at 21 dpi were classified by severity of infection (low versus high lesions), we wanted to know if host gene expression was different between these two groups. Therefore, animals with high level of lesions were compared to those with low or minimal lesions. With this comparison, there were many more DEGs (n = 494) observed than when comparing all infected animals together to non-infected animals (n=22). At 28 dpi the comparison between high and low lesion animals is different than that of 21dpi because two of the three low lesion animals had either a negative HE or IHC score and did not shed *L. intracellularis* at this time point, suggesting they had cleared infection or had a minimal level of infection. At 28 dpi there were 23 DEG comparing animals with high level lesions to those with low or no lesion levels. Figure 2 shows a Venn diagram comparing the number of DEGs between groups of high and low lesion animals and infected and non-infected animals at different time points.

**Figure 2.**
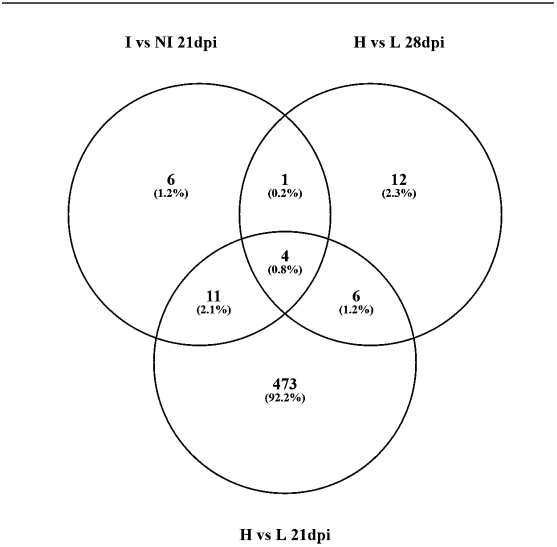
Venn diagram of differentially expressed genes compared between infected and non-infected animals at 21 dpi (I vs NI 21dpi), high and low lesion animals at 21 dpi (H vs L 21dpi) and high and low lesion animals at 28 dpi (H vs L 28dpi).

We focused on the time point of 21dpi because this is the time point when peak of infection occurred as measured by fecal shedding and the number of animals with lesions and their severity. The ten most upregulated and downregulated genes identified when comparing animals with high lesions to those with low lesions at 21 dpi are shown in table 3. Similar to the DEG identified comparing challenged to non-challenged animals at 21dpi, *MMP7, GABRP* and *SERPINB2* were also among the most differentially expressed genes. Other upregulated genes included: adhesion G protein-coupled receptor A2 (*ADGRA2),* bradykinin receptor B1 (*BDKRB1),* TNF Alpha Induced Protein 6 (*TNFAIP6*) and integrin subunit alpha 8 (*ITGA8*). Among the most downregulated genes in high compared to low lesion animals at 21 dpi were two genes that encode the apolipoproteins C-III (*APOC3*) and A-IV(APOA4) as well as the gene that encodes fibroblast growth factor 19 protein (*FGF19*). These code for proteins which are expressed in the small intestine and play important roles in lipid metabolism among other functions (21–23).

**Table 3.**
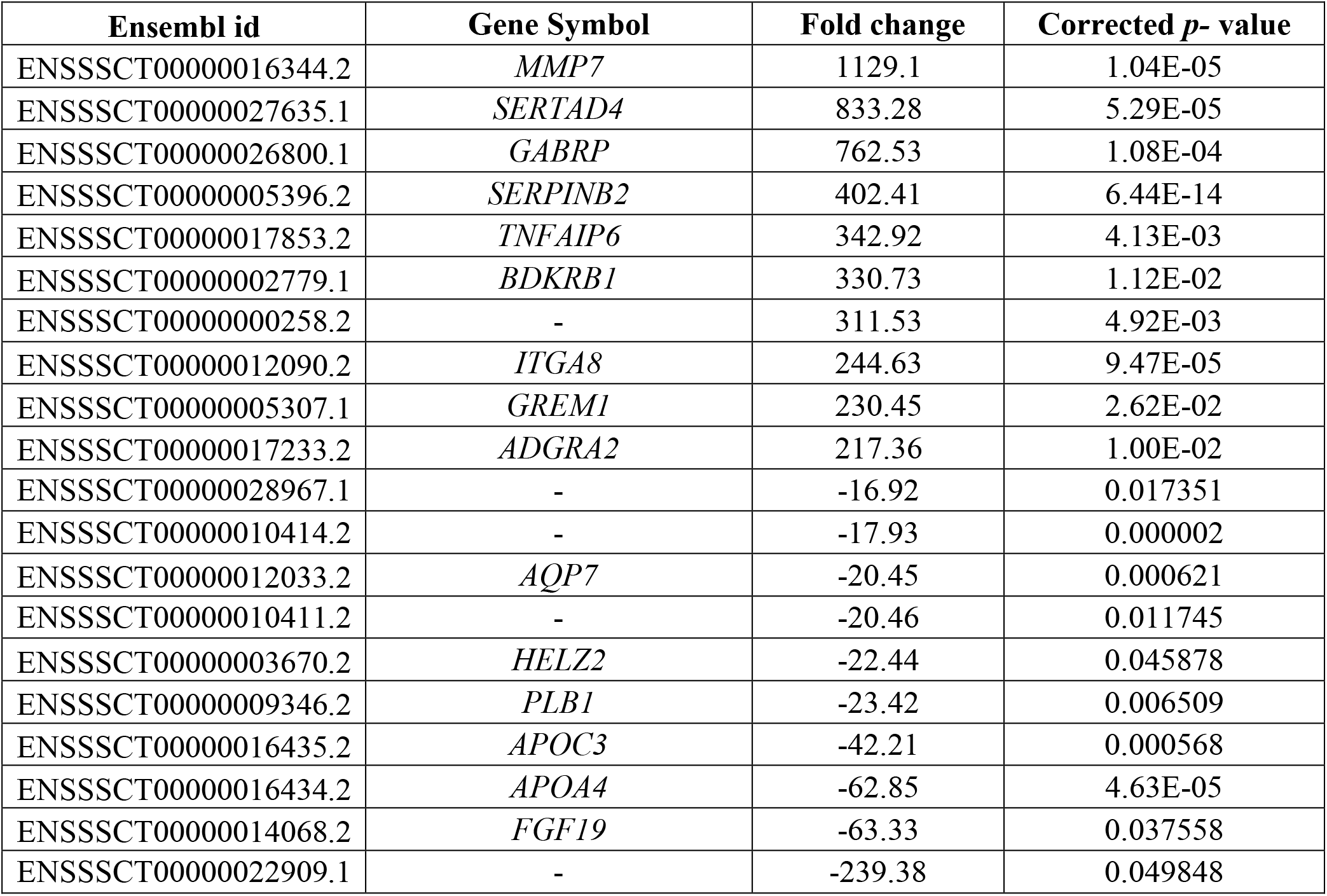
The ten most upregulated (among the 417 upregulated) genes and the ten most downregulated (among the 77 downregulated) genes at 21 dpi comparing animals with high and low lesions.

Among the 23 genes differentially expressed between high and low lesion animals at 28 dpi (table 4), four were in common between infected and non-infected animals as well as between high and low lesion animals at 21 dpi (Figure 2). These genes were *XDH, MMP7, TGM2* and a porcine gene with no known human or mouse homologue (ENSSSCT00000014604.2). *IDO1,* which encodes indoleamine 2 3-dioxygenase, was up regulated in infected animals at 21 dpi as well as in high lesion animals at 28dpi. The gene *MGP,* as well as the genes *TMEPAI* and *HTRA3,* which have been found to play roles in apoptosis and influencing cell proliferation, were up regulated in high lesion animals of both 21 and 28 dpi (24, 25). Around half of the 23 DEG of high lesion animals at 28dpi were only found at that time point. Several of these genes encode proteins involved in antibody biosynthesis, including *IGLC1* that encodes the constant region of immunoglobulins and *IGLV-10,* and *IGLV-12* that encode the variable regions of the lambda light chain of immunoglobulins (26). The only gene down regulated in animals with high lesions at 28 dpi was *CA1* which codes for carbonic anhydrase 1 (27).

**Table 4.**
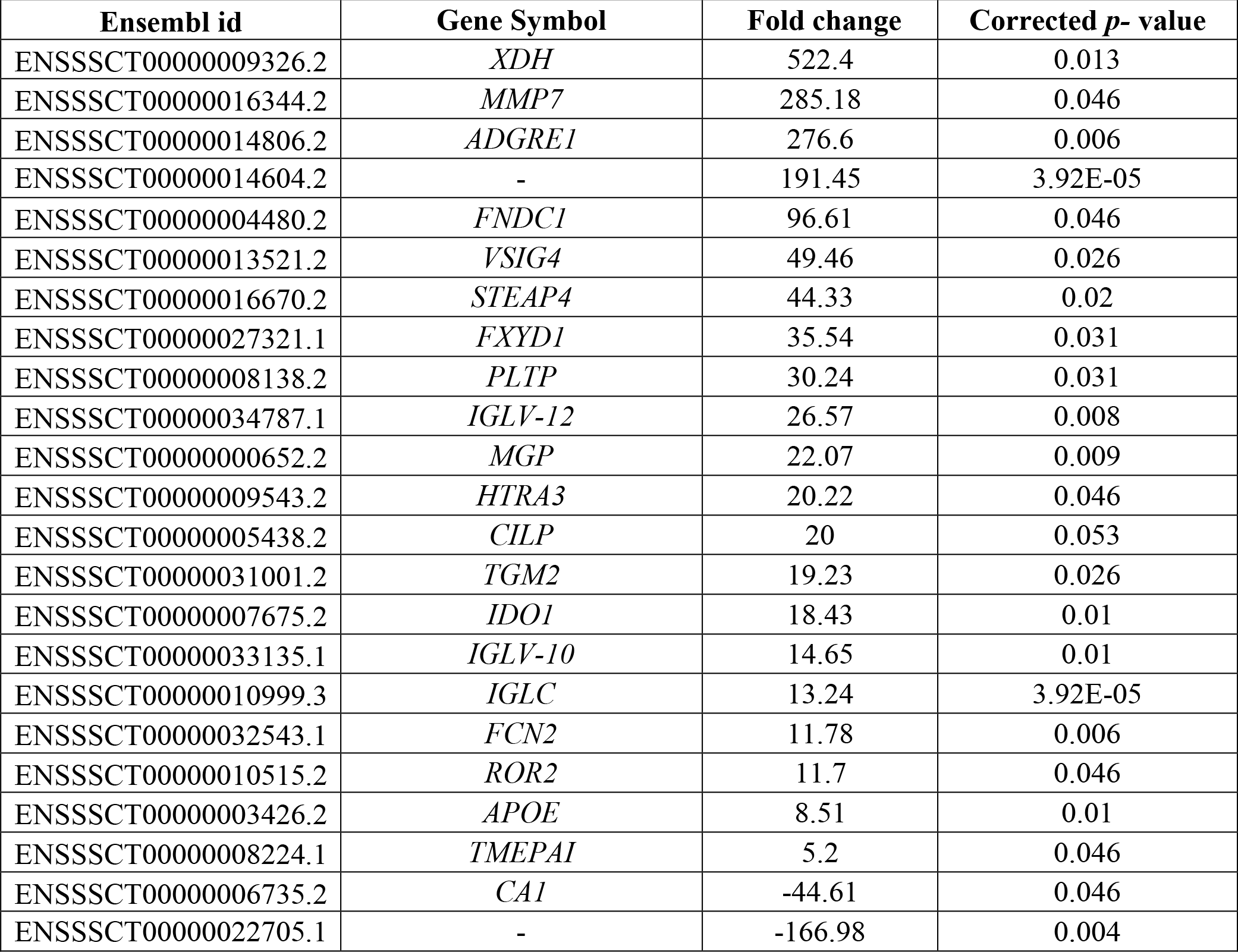
Differentially expressed genes between high and low lesion animals at 28 days post infection.

### Pathway Analysis: biological functions associated with increased infection

To explore potential biological functions associated with the hyperplasia that is characteristic of *L. intracellularis* infection, biological functions associated with “proliferation” were investigated. This analysis revealed 16 biological functions significantly associated (*p*<0.05) with the DEG and the most significant was “cell proliferation of tumor cell lines” (*p* = 1. 4E-15, table 5). IPA analysis attributes an activation z-score that assesses the randomness of directionality within a gene set to infer the activation state of a metric (i.e. biological function, upstream regulator or pathway) as either activated or inhibited. A negative z-score indicates inhibition and a positive z-score indicates activation with scores equal to or greater than 2 or equal to or less than −2 being statistically significant (*p*<0.05) (28). Of the 16 biological functions associated with proliferation, six had significant z-scores above 2 including “cell proliferation of tumor cell lines” (table 5).

**Table 5.** List of diseases and functions associated with “proliferation” among the diseases and functions identified between animals with high and low lesions using Pathway Analysis.

| **Function Annotation** | ***p*-value** | **z-score** | **Number of Molecules** |
| --- | --- | --- | --- |
| proliferation of connective tissue cells | 4.75E-14 | 3.983 | 50 |
| cell proliferation of fibroblasts | 4.32E-09 | 3.873 | 29 |
| proliferation of synovial cells | 5.92E-08 | 2.922 | 9 |
| cell proliferation of tumor cell lines | 1.4E-15 | 2.671 | 98 |
| proliferation of stem cells | 0.000014 | 2.52 | 16 |
| cell proliferation of breast cancer cell lines | 0.000000235 | 2.075 | 32 |
| proliferation of pericytes | 0.0000125 | 1.71 | 8 |
| endothelial cell development | 0.000000131 | 1.461 | 27 |
| proliferation of liver cells | 0.0000153 | 1.25 | 15 |
| proliferation of tumor cells | 0.0000106 | 1.22 | 28 |
| proliferation of endothelial cells | 9.05E-08 | 1.142 | 25 |
| proliferation of epithelial cells | 7.77E-09 | 1.007 | 36 |
| cell proliferation of ovarian cancer cell lines | 0.00000357 | 0.891 | 13 |
| colony formation of cells | 4.52E-08 | 0.429 | 34 |
| colony formation | 1.85E-08 | 0.401 | 37 |
| colony formation of tumor cell lines | 0.00000122 | -0.537 | 22 |

### Pathway Analysis: canonical pathways associated with increased infection

Next, we used IPA to identify canonical pathways associated with the DEGs between high and low lesion animals at 21 dpi. Of these pathways, we searched for pathways with a significant activation status. This analysis revealed 10 canonical pathways significantly activated associated with the data set (z-score of ≥ 2, table 6).

**Table 6.**
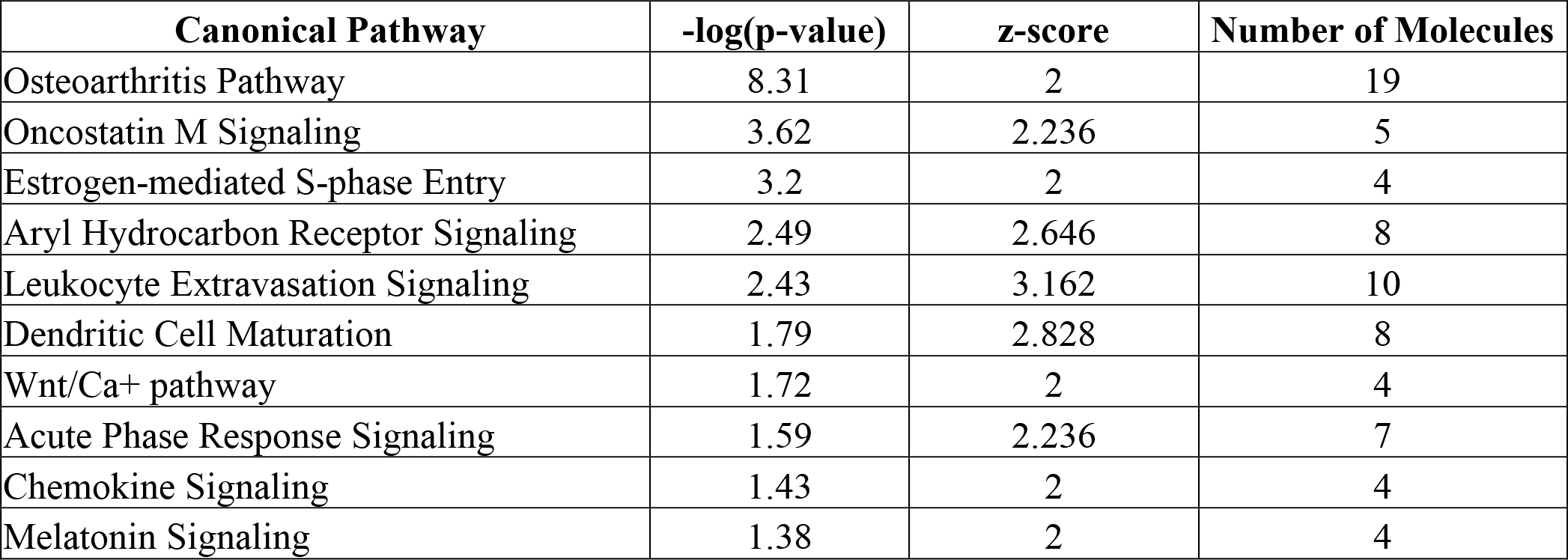
Activated canonical pathways in animals with high lesions compared to low lesions animals at 21 days post infection.

| **Canonical Pathway** | **-log(p-value)** | **z-score** | **Number of Molecules** |
| --- | --- | --- | --- |
| Osteoarthritis Pathway | 8.31 | 2 | 19 |
| Oncostatin M Signaling | 3.62 | 2.236 | 5 |
| Estrogen-mediated S-phase Entry | 3.2 | 2 | 4 |
| Aryl Hydrocarbon Receptor Signaling | 2.49 | 2.646 | 8 |
| Leukocyte Extravasation Signaling | 2.43 | 3.162 | 10 |
| Dendritic Cell Maturation | 1.79 | 2.828 | 8 |
| Wnt/Ca<sup>+</sup> pathway | 1.72 | 2 | 4 |
| Acute Phase Response Signaling | 1.59 | 2.236 | 7 |
| Chemokine Signaling | 1.43 | 2 | 4 |
| Melatonin Signaling | 1.38 | 2 | 4 |

### Pathway Analysis: upstream regulators of differentially expressed genes

Potential regulator molecules responsible for the differential gene expression of the different pathways observed at 21dpi were identified using IPA. Upstream regulators are molecules that are known to influence the observed DEG and are potentially responsible for modulating their differential expression (28). Cyclin dependent kinase inhibitor 1A (CDKN1A, p = 2.06E-36), tumor protein p53 (TP53, p = 9.48E-29) and Erb-b2 receptor tyrosine kinase 2 (ERBB2, *p* = 2.57E-26) were among the upstream regulators most significantly associated with the dataset. Other significant upstream regulators included the cytokines transforming growth factor beta 1 (TGFB1, *p* = 3.51E-28) tumor necrosis factor (TNF, *p* = 2.23E-23), IL-1β and IL-6 (table 7).

**Table 7.** The 30 most significant up-stream regulators associated with the differentially expressed genes between high and low infected animals.

| **Upstream Regulator** | **Fold Change** | **Molecule Type** | **z-score** | **p-value** |
| --- | --- | --- | --- | --- |
| CDKN1A |  | kinase | -2.696 | 2.06E-36 |
| TP53 |  | transcription regulator | -1.797 | 9.48E-29 |
| TGFB1 |  | growth factor | 4.163 | 3.51E-28 |
| ERBB2 |  | kinase | 4.368 | 2.57E-26 |
| TNF |  | cytokine | 2.31 | 2.23E-23 |
| Calcitriol |  | chemical drug | -2.059 | 1.12E-20 |
| HGF |  | growth factor | 3.546 | 1.94E-20 |
| RABL6 |  | other | 4.472 | 4.4E-20 |
| lipopolysaccharide |  | chemical drug | 4.252 | 1.22E-19 |
| CCND1 |  | transcription regulator | 2.339 | 2.05E-19 |
| beta-estradiol |  | chemical - endogenous mammalian | 3.237 | 4.04E-19 |
| Tretinoin |  | chemical - endogenous mammalian | 2.096 | 6.88E-19 |
| Vegf |  | group | 4.611 | 1.04E-18 |
| PDGF BB |  | complex | 3.059 | 1.07E-18 |
| IL6 |  | cytokine | 3.836 | 1.88E-18 |
| dextran sulfate |  | chemical drug | 1.627 | 3.7E-18 |
| SMAD7 |  | transcription regulator | -1.759 | 5.52E-18 |
| medroxyprogesterone acetate |  | chemical drug | -2.449 | 5.52E-18 |
| E2F4 |  | transcription regulator | -0.927 | 3.52E-17 |
| Alpha catenin |  | group | -4.145 | 9.43E-17 |
| PTGER2 |  | g-protein coupled receptor | 3.702 | 1.32E-16 |
| IL1B | 29.3 | cytokine | 3.216 | 1.53E-16 |
| Tgf beta |  | group | 1.365 | 3.53E-16 |
| cycloheximide |  | chemical reagent | 0.371 | 5.29E-16 |
| dexamethasone |  | chemical drug | 0.526 | 1.41E-15 |
| EP400 |  | other | 3.148 | 2.43E-15 |
| CDKN2A |  | transcription regulator | -2.69 | 5.15E-15 |
| butyric acid |  | chemical - endogenous mammalian | -0.573 | 8.37E-15 |
| trichostatin A |  | chemical drug | 1.112 | 1.77E-14 |
| MYC | 4.2 | transcription regulator | 2.574 | 2.78E-14 |

### Matrix metalloproteinase-7 immunohistochemistry

To determine if changes observed at the level of transcription of *MMP7* were translated into protein in infected tissue, we chose to perform immunohistochemistry staining for matrix metalloproteinase-7 (MMP-7). We observed that the distribution of staining of this molecule differed between infected and non-infected animals. Staining of MMP-7 was not associated with hyperplastic crypts. MMP-7 positive cells were observed in normal crypts adjacent to those undergoing hyperplasia. Staining was observed both in the apical side of enterocytes as well as associated with the lumen of crypts (Figure 3). This pattern of staining was not observed in non-infected animals (data not shown).

**Figure 3.**
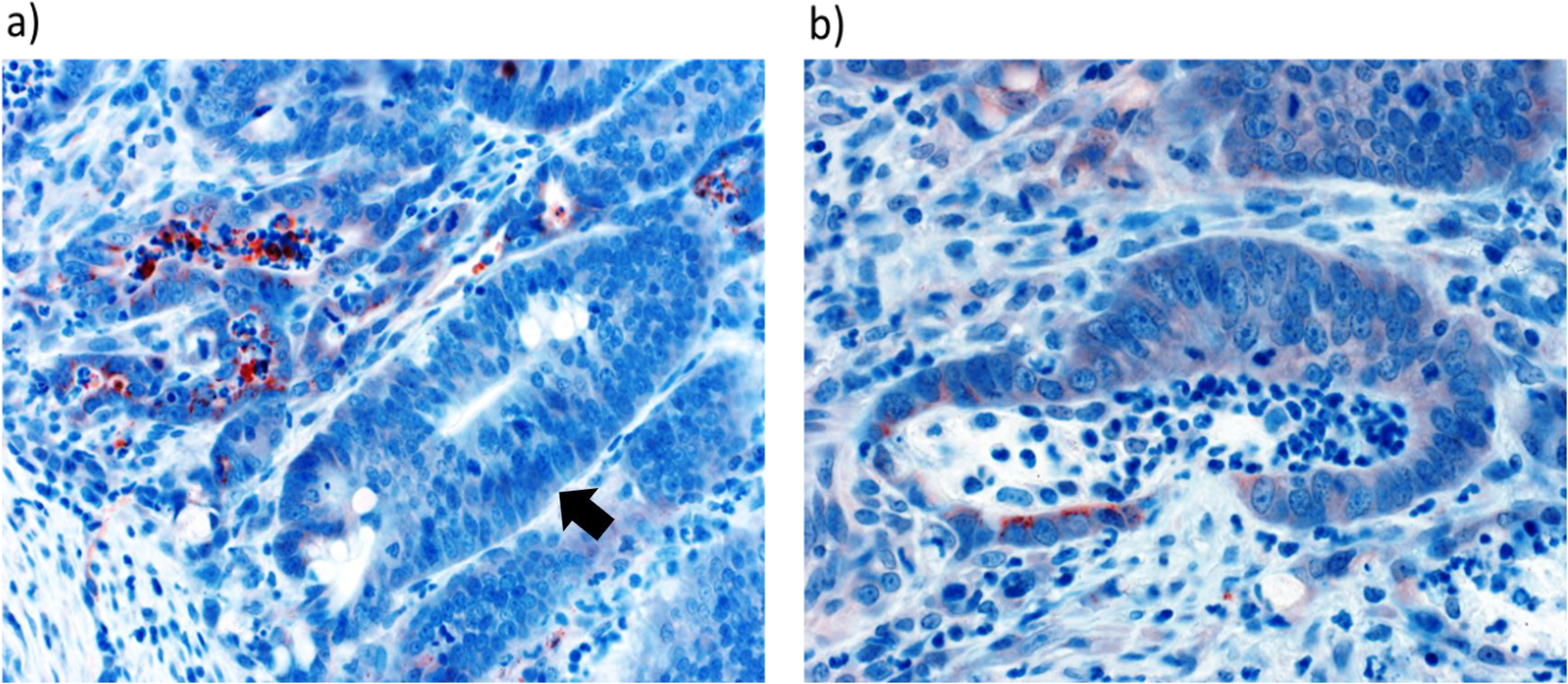
Matrix metalloproteinase-7 (MMP-7) staining. a) Luminal staining of MMP-7 (red) in crypts adjacent to a hyperplastic crypt (arrow), b) apical staining of MMP-7 (red) enterocytes in crypt.

## Discussion

In this study we reproduced PPE with the induction of characteristic microscopic and macroscopic lesions. We also assessed clinical signs and as is most cases of PIA in the field, animals had subclinical disease (data not shown). Infection was also confirmed by quantifying *L. intracellularis* shed in feces and the presence of antibodies in serum. We observed a time effect of disease that has previously been described in which hyperplasic lesions begin at around 11 dpi, peak at around 21 dpi and begin to resolve at 28 dpi (29). Changes to crypt depth, fecal shedding and villus height were observed and were more severe at 21dpi in infected animals. This is in accordance with previous studies and suggests that peak of infection occurred at this time point. At 28 dpi two out of the six animals had either negative IHC or HE scores and did not shed *L. intracellularis,* indicating a likely resolution of infection, as is known to occur in this disease (1). At 28 dpi, animals also showed high antibody titers against *L. intracellularis* indicative of a strong prior immune response to infection.

In this study, at 14 dpi the lack of differentially expressed genes coincided with low lesion severity and antigen load. This was also observed by Smith et al. 2014 who analyzed *L. intracellularis* infected intestine tissue samples at different time points by microarray analysis and found that the number of DEG coincided with the quantity of *L. intracellularis* present in tissue. This indicates there isn’t a robust host response during early stages of the infection and this coincides with lower lesions. At 21 dpi, when animals developed severe lesions, there were 22 DEG between infected and non-infected animals. When comparing animals with high to low level of infection and lesions at this time point there were 494 DEG, again suggesting that development of severe lesions is associated with a greater host response. Several of the genes found to be differentially expressed in this study were not observed by Smith et al., 2014, although “cancer” was the major disease category found with pathway analysis in both studies (data not shown, *p* = 5.86E-23-1.9E-05). Differences in gene expression between both studies may be due to differences in lesion severity caused by infection and/or due to differences between microarray and RNAseq methods of gene expression analysis.

The gene most highly and differentially expressed in infected animals at 21 dpi was *XDH* (816.98 fold higher in infected animals, *p* = 0.009, table 2) which encodes xanthine dehydrogenase. This gene was also differentially expressed between high and low lesions animals at both 21 and 28 dpi. Xanthine dehydrogenase is a key regulator of inflammatory cascades whose products include reactive oxygen and nitrogen species (30, 31). This enzyme is highly expressed in the small intestine and has anti-microbial properties (31). Recently, it was found to be highly expressed in *Mycobacterium avium* subspecies *paratuberculosis* infection where it was suggested to be indicative of macrophage activation and killing of the pathogen. A similar response is possible in infections with *L. intracellularis* (32).

*MMP7,* which encodes matrix metalloproteinase-7 or matrilysin, was also a gene that was found to be up regulated in infected as well as in animals with high lesion scores at 21 and 28 dpi. This protein can be induced by some gram-negative bacteria and it has an important role in activating α-defensin antimicrobial peptides (33, 34). Pathogenic strains of *Helicobacter pylori,* which have a greater capacity to cause gastric cancer, induce *MMP7* expression in a cag pathogenicity island dependent manner (35, 36). Overexpression of *MMP7* is commonly found in colorectal cancers and this protein can induce increased cellular proliferation (37–40). Interestingly, positive staining of this protein in ileal tissues from animals with high lesion was found in crypts adjacent to those undergoing hyperplasia (Figure 3).

Within the diseases and functions associated with the DEG data-set involving proliferation, the biological function of “cell proliferation of tumor cell lines” was the most significantly associated with the dataset and had a significant activated status (*p* = 1.4E-15, z-score of 2.67). This function had the involvement of 98 of the 494 DEG among animals with high lesion scores at 21 dpi, which also correlated with the time when peak of crypt depth was noted using histomorphometry (Figure 1a). The genes *MMP7, OSM, SERPINB2, GABRP, TACSTD2* and *TGM2* differentially expressed between infected and non-infected animals at 21 dpi were also among these 98. This demonstrates that the molecular signature induced by *L. intracellularis* is strongly associated with cellular proliferation, several molecules of which have also been found in to be up-regulated in different tumor cell lines. This makes sense since hyperplasia of enterocytes is observed in *L. intracellularis* infection. The two upstream regulators most significantly associated with the data set, cyclin dependent kinase inhibitor 1A (CDKN1A, p = 2.06E-36) and tumor protein p53 (TP53, p = 9.48E-29), are also associated with cell proliferation. CDKN1A inhibits the cyclin kinases that are necessary for cell cycle progression during the G1 and S phases and has an antiproliferative effector function. This gene is induced by p53 tumor suppressor protein which is activated upon cellular stressors to induce the expression of genes that result in inhibition of proliferation (41, 42). Both of these regulators had negative z-scores indicating that they were down regulated, thus likely not maintaining their function of inhibiting cell proliferation (table 7).

There is a large body of evidence to suggest that tumor-associated calcium signal transducer 2 (*TACSTD2,* also named Trop2) promotes cellular proliferation and this gene is highly expressed in a wide variety of epithelial tumors (43, 44). Interestingly, it has also been found that knocking down of this gene in mice can promote carcinogenesis in a squamous cell cancer model, which also suggests a protective role of this protein (45). Similarly, transglulatminase-2 (TGM2) can also promote cell proliferation although this protein has many other functions (46). *TGM2* expression is induced by inflammatory mediators such as IL-1β, TNFα and TGF-β and it has been described to be involved in inflammatory conditions of the intestine (20, 47). TNF, IL-1β and TGF-β were found to be upstream regulators (table 7) and are likely contributing to the host response to *L. intracellularis*. Interestingly, a common histomorphological feature of inflammatory conditions of the intestine is hyperplasia of enterocytes which are marked by an increase in epithelial cell numbers that are visible as elongated crypts (48). Although not statistically significant, we found that animals with high lesions had greater average crypt depth demonstrating that crypts are enlarged with more severe infection (Figure 1a) and elongated crypts are a common feature of PPE (1). Two other histomorphological features of intestinal inflammation are villus blunting and loss of goblet cells (48, 49). We also observed a decrease in villus height with infection (Figure 1b), and the decrease or absence of goblet cells is known to occur in *L. intracellularis* infection (1). This suggests that the host response to infection may contribute to the altered state of enterocytes in addition to any potential mechanism *L. intracellularis* may have itself.

Also associated with inflammation is the gene *OSM* which encodes Oncostatin M, a member of the IL-6 cytokine family, expressed by activated T cells, monocytes, and dendritic cells (18). Increased expression of this cytokine has been found in inflammatory bowel disease, where it correlated to histopathological disease severity (50, 51). OSM is also capable of inducing enterocyte proliferation (50). In our data set not only was the *OSM* gene up-regulated with infection but its canonical pathway was also significantly associated and activated in animals with high lesion scores at 21 dpi (table 6).

Several of the other canonical pathways and differentially expressed genes were also involved in inflammation and immune response. These include the pathways of leukocyte extravasation signaling, dendritic cell maturation and acute phase response signaling. Increased levels of acute phase proteins have previously been found in pigs infected with *L. intracellularis* (52). This gives some insight into the mucosal immune response to infection which is known to be protective and suggests that antigen presentation with dendritic cells and the presence of leukocytes are an important part of this process. Aryl hydrocarbon receptor (AhR) signaling is becoming increasingly recognized as an important pathway that influences the immune responses to pathogens (53). This pathway was significantly activated in animals with high lesion at 21 dpi (table 6). The precise function of this signaling pathway varies with different pathogens and affected organs (53). In infections of mice with *Citrobacter rodentium,* a bacterial pathogen that also infects the gut and induces hyperplasia of enterocytes (54), knockdown of the *AhR* led to significant weight loss and death while control animals did not succumb to infection (55). Perhaps activation of AhR signaling at the peak of infection is protective to *L. intracellularis* infection and contributes to the resolution of lesions with time.

Investigating the genes that were differentially expressed exclusively at 28 dpi gives some insight into the host response associated with the start of resolving lesions. The only gene differentially expressed between infected and non-infected animals at this timepoint was *UNC5B* (unc-5 netrin receptor B), with a 3.77 fold higher expression in infected animals (*p* < 0.05). This gene encodes a transmembrane receptor that can regulate cell fate as a “dependence receptor” and induces apoptosis in the absence of its ligand (56, 57). Higher expression of *UNC5B* can inhibit cellular proliferation in some cancer cell lines and this protein is inactive and down regulated in colorectal cancer (56, 58, 59). Additionally, increased expression of *UNC5B* in leukocytes has been shown to attenuate inflammation (60). Thus higher expression of *UNC5B* may be involved in inhibiting enterocyte hyperplasia and/or resolving inflammation. The only gene found to be expressed higher in low lesions animals compared to high lesions animals at this time point was *CA1* (carbonic anhydrase 1, CA-1). Carbonic anhydrases can be found along the entire gastro-intestinal tract, and in the small intestine carbonic anhydrase 1 is found in cryptal enterocytes (61). Interestingly, increased expression and staining of *CA-1* is found in healthy intestinal mucosa where it participates in regulation of pH homeostasis and water and ion transport while decreased CA-1 protein and mRNA levels can be found in benign and malignant colorectal tumors (61–63). Lastly, several of the genes only found in animals with high lesion at 28 dpi were associated with synthesis of immunoglobulins. This gives important insight as to the active immune response that is occurring at this stage of infection.

The research described here leads us to conclude that the host response to infection leads to the induction of several genes and pathways associated with cellular proliferation with some similarity to what has been described in some cancers and inflammatory diseases of the intestinal tract. However, infections caused by *L. intracellularis* are very much different from cancer as the proliferative lesions do not develop into tumors and in fact resolve. The mechanisms that lead to the expression of the observed genes associated with increased cellular proliferation should be investigated further as well as their specific contribution to disease. The return to a normal state of the intestine without the expression of these genes is likely due to a protective immune response that clears the pathogen. This research sheds important light into the pathogenesis of *L. intracellularis* demonstrating several of the genes and pathways associated with its pathology.

## Acknowledgements

We would like to acknowledge the Zinpro corporation who funded a part of this project. We would also like to thank Jan Shivers and Molly Uaje for performing staining of histological slides. This work was funded by a USDA Multistate Health Formula Fund. The RNA seq data set is available at the following site: https://doi.org/10.13020/D63M5F.

